# The early human interferon gamma response to *Toxoplasma gondii* is driven by Vγ9Vδ2 T-cell sensing of host phosphoantigens and subsequent NK-cell activation

**DOI:** 10.1101/2025.06.12.659293

**Authors:** Felipe Rodriguez, Jeroen P.J. Saeij

**Affiliations:** Department of Pathology, Microbiology and Immunology, School of Veterinary Medicine, University of California, Davis, CA 95616

## Abstract

*Toxoplasma gondii* is a globally prevalent intracellular parasite that infects ∼40 million Americans. The murine immune response to *Toxoplasma* relies on both toll-like receptor (TLR) 11/12 and immunity related GTPase-mediated (IRGs) responses, which humans lack, making it unclear how the human immune response detects and responds to the parasite. We investigated whether human Vγ9Vδ2 T cells, which detect phosphoantigens through the BTN3A1 receptor, shape the early immune response to the parasite. Using primary human peripheral blood mononuclear cells (PBMCs), we show that Vγ9Vδ2 T cells are activated by *Toxoplasma*-infected cells in a BTN3A1- dependent manner leading to secretion of interferon gamma (IFNγ) and tumor necrosis factor-alpha (TNFα). Additionally, these T cells potentiate IFNγ production by natural killer (NK) cells, likely via TNFα and interleukin (IL)-12 produced during infection. Active parasite invasion is required to stimulate the IFNγ response, and inhibition of the host mevalonate pathway, which limits the synthesis of the phosphoantigen isopentenyl pyrophosphate (IPP), attenuates the cytokine response, indicating *Toxoplasma* infection increases host phosphoantigens leading to Vγ9Vδ2 T cell activation. Our findings identify Vγ9Vδ2 T cells as key effectors that potentiate NK cells in the early human immune response to *Toxoplasma*, bridging innate and adaptive immunity in the absence of TLR11/12 signaling.

## Introduction

*Toxoplasma gondii* is an intracellular apicomplexan parasite that remains a leading cause of foodborne mortality. Felines are its definitive host and can shed millions of infectious oocysts, leading to widespread environmental contamination (1,2). Following ingestion of oocysts or tissue cysts, the parasite disseminates and, by converting to slowly replicating bradyzoites within tissue cysts, establishes long-lived infection. This form is notably resistant to immune clearance, and reactivation of tissue cysts in immunosuppressed individuals can cause severe pathology including brain lesions and necrotizing retinitis, potentially resulting in death. *Toxoplasma* can also cause miscarriage in pregnant mothers infected for the first time (3). Inside host cells, *Toxoplasma* resides in a parasitophorous vacuole (PV), separated from the host cell cytosol by the PV membrane (PVM), which offers a replication niche (4).

In mice, toll-like-receptor (TLR) 11/12 activation by *Toxoplasma* actin-binding protein profilin, triggers dendritic cells (DCs) to secrete interleukin (IL)-12 which activates natural killer (NK), CD4, and CD8 T cells to secrete interferon gamma (IFNγ), which can be further enhanced by IL-18 and IL-1β (5–7). IFNγ upregulates multiple interferon-stimulated genes with parasiticidal properties. In mice, IFNγ upregulates the expression of immunity-related GTPases (IRGs), which can destroy the PV and thereby eliminate *Toxoplasma*’s replication niche. *Toxoplasma* secretes a plethora of effector proteins (dense granule antigens (GRAs) and rhoptry bulb proteins (ROPs)) into the host cytosol that hijack host cell signaling pathways and prevent the host from mounting an appropriate immune response (8,9). In mice, GRAs and ROPs block IRGs from destroying the PV (10). However, humans lack TLR11/12 and IFNγ-inducible IRGs, and most of the *Toxoplasma* ROPs and GRAs that determine virulence in rodents play no role in human infection. Thus, we currently have an incomplete understanding of what human immune cells are orchestrating cytokine secretion to *Toxoplasma*, and how the human immune system detects and destroys *Toxoplasma*.

Early studies have shown that *Toxoplasma* can infect any nucleated cell and use blood to spread throughout the host (11,12) Human peripheral blood mononuclear cells (PBMCs) consist of ∼10% monocytes, ∼60% T cells, and ∼15% NK cells (13), which play an important role in defense against infections. Vγ9Vδ2 T cells, which constitute up to 3% of human peripheral T lymphocytes and dominate the circulating γδ pool, recognize non-peptidic prenyl-pyrophosphate antigens such as (E)-4-hydroxy-3-methyl-but-2-enyl pyrophosphate (HMBPP) and isopentenyl pyrophosphate (IPP) derived from bacteria such as *Mycobacterium tuberculosis* (Mtb) and *Listeria monocytogenes* or protozoan parasites like *Plasmodium* spp. (14–18). Additionally, tumor cells upregulate IPP via the mevalonate pathway, which can stimulate Vγ9Vδ2 T cells leading to cell death (19). Phosphoantigen-reactive Vγ9Vδ2 T cells exist only in humans, non-human primates, and alpacas and make up to 90% of total peripheral blood γδ T cells in healthy adults (20,21). Vγ9Vδ2 T cells are important in both innate and adaptive immunity and mount major expansion and effector responses during infections (22–24). Vγ9Vδ2 T cells do not detect phosphoantigens directly but instead bind to the butyrophilin 2A and 3A (BTN2A1/BTN3A1) dual receptors. Phosphoantigens bind to the intracellular B30.2 domain of the BTN3A1 receptor promoting the hetero-dimerization to BTN2A1, which is subsequently detected by Vγ9Vδ2 T cells resulting in their activation and secretion of IFNγ and TNFα, leading to activation of host cells (19,25,26).

*Toxoplasma* utilizes a specialized organelle known as the apicoplast, which evolutionarily is of bacterial origin, for synthesizing isoprenoid precursors through the methyl-erythritol phosphate (DOXP) pathway, producing HMBPP and theoretically providing a ligand for Vγ9Vδ2 T cell activation (27). Limited observations suggest Vγ9Vδ2 T cells are cytotoxic toward *Toxoplasma*-infected cells (28) and increase numbers in congenitally infected fetuses (29), but a mechanistic link between parasite-derived HMBPP, butyrophilin receptor engagement, and early IFNγ release has not been demonstrated.

Here we show that *Toxoplasma* indirectly activates human Vγ9Vδ2 T cells via BTN3A1, leading to rapid IFNγ secretion and potentiation of NK cells. These findings support a model in which Vγ9Vδ2 T cells compensate for the absence of TLR11/12 in humans by providing an early source of IFNγ and by licensing other innate lymphocytes, thereby constraining parasite replication before conventional αβ T-cell immunity is established.

### BTN3A1-dependent IFNγ production by human Vγ9Vδ2 T cells in response to HMBPP

Considering that PBMCs contain a diverse repertoire of T and NK cells, it remains unclear which specific cell types are driving cytokine production during *Toxoplasma* infection. Given that *Toxoplasma* utilizes the 1-deoxy-D-xylulose-5-phosphate (DOXP) pathway for lipid and HMBPP production (27,30), we hypothesized that Vγ9Vδ2 T cells are activated upon *Toxoplasma*-derived HMBPP exposure.

To optimize our protocols, we first exposed PBMCs from *Toxoplasma*-seronegative donors to commercially sourced HMBPP (Fig S1A), either alone or with apyrase, a terminal phosphatase that can cleave HMBPP, thus preventing its stimulatory effect on Vγ9Vδ2 T cells via BTN3A1/BTN2A1 receptors (31,32). Exogenous HMBPP significantly promoted IFNγ secretion from human PBMCs, which was abrogated by apyrase (Fig 1A). Because Vγ9Vδ2 T cells recognize phosphoantigens via BTN3A1, we next tested whether blocking this receptor affects the response. Addition of a BTN3A1 blocking antibody, but not an IgG1 isotype control antibody, blocked IFNγ secretion, showing that IFNγ secretion was dependent on the BTN3A1 receptor (Fig 1A). Flow cytometry was then used to investigate the source of IFNγ in response to HMBPP. The gating strategy is shown in (Fig S1B). Baseline frequencies of Vγ9Vδ2 T cells differed between donors, likely explaining some of the variability in our assays (Fig 1B). HMBPP significantly increased the percentage of IFNγ-producing CD3^+^Vγ9^+^ cells, which was inhibited by both apyrase and BTN3A1 blocking antibody. Despite inter-donor variability, the percentage of IFNγ-positive Vγ9^+^ T cells was significantly reduced when apyrase or BTN3A1 blocking antibody were added with HMBPP (Fig 1C). Integrated mean fluorescent intensity (iMFI) analysis showed that iMFI levels in IFNγ-positive Vγ9Vδ2 cells were significantly higher in HMBPP-stimulated cells compared to those where HMBPP-stimulated cells were also treated with either apyrase or BTN3A1 blocking antibody (Fig 1D) (33). To determine whether HMBPP activates other immune cells to secrete IFNγ, we gated IFNγ-positive CD3^+^Vγ9^-^ (Fig 1E), and IFNγ-positive CD3^-^ (Fig 1F), which encompasses CD4 and CD8 T cells, and NK cells, respectively, and noticed no difference in percentage of IFNγ^+^ cells. Overall, these results demonstrate that 1) donors vary in Vγ9^+^ T cell frequency in blood, 2) these cells are the primary mediators of IFNγ secretion when stimulated with exogenous HMBPP, and 3) HMBPP-mediated induction of IFNγ is blocked both by apyrase and BTN3A1 blocking antibody.

**Fig 1.**
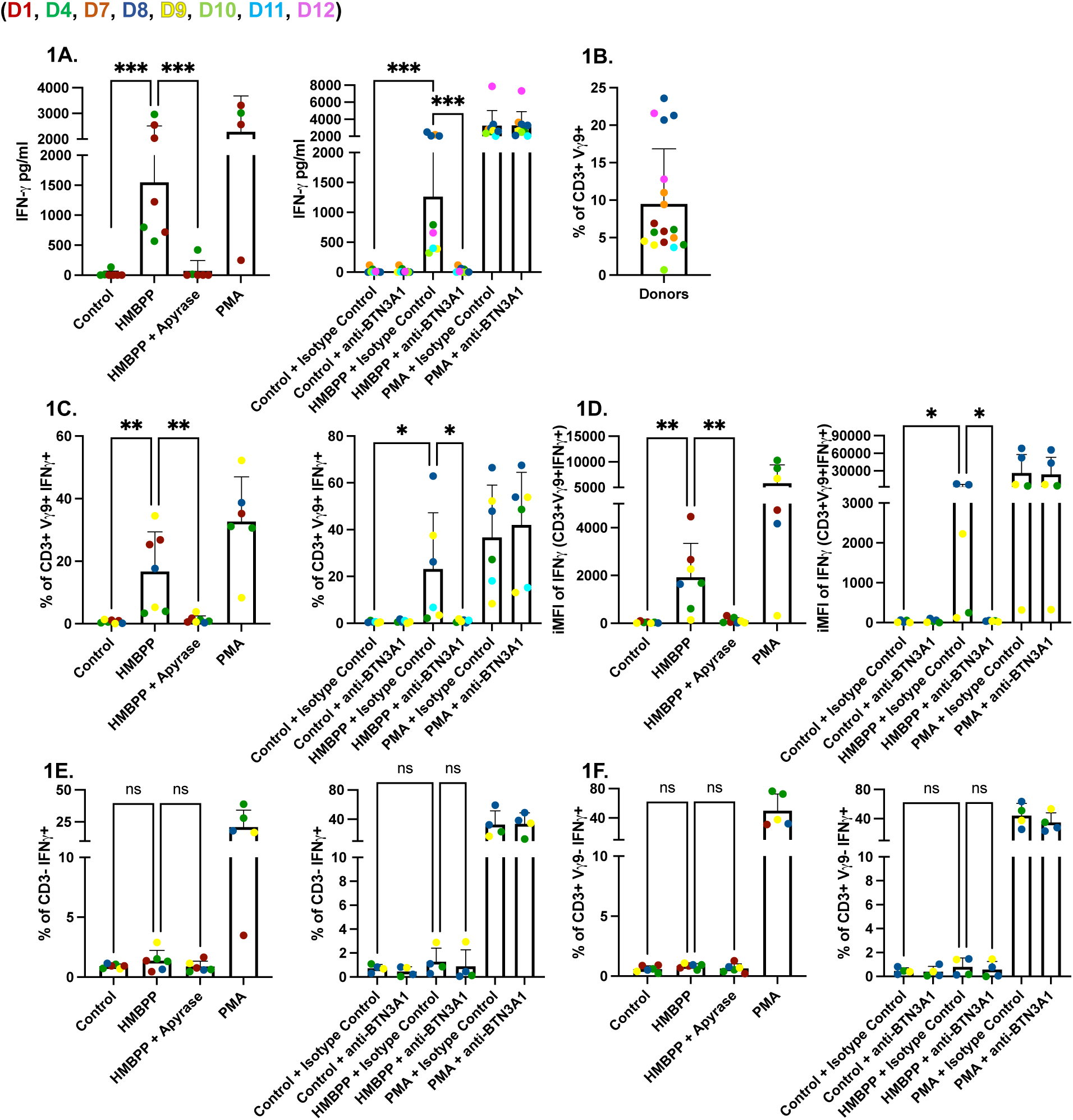
HMBPP stimulates human Vγ9Vδ2 T cells leading to the production of IFNγ. **(A)** PBMCs (5×10^5^/well) were left untreated or were incubated for 24 hours with HMBPP (312 nM) or PMA/ionomycin (1x). Cultures contained either apyrase (200 IU/ml), or BTN3A1 blocking antibody (0.1 µg/ml), or IgG1 isotype control antibody (0.1 µg/ml). IFNγ concentrations (pg/ml) in culture supernatants were measured by ELISA. **(B)** The percentage of CD3^+^Vγ9^+^ cells in each donor. **(C)** Percentages of lymphocytes/single cells/alive/CD3^+^/Vγ9^+^/IFNγ^+^. **(D)** (iMFI) of IFNγ in Vγ9^+^. **(E)** Percentages of lymphocytes/single cells/alive/CD3^-^/IFNγ^+^. **(F)** Percentages of lymphocytes/single cells/alive/CD3^+^/Vγ9^-^/IFNγ^+^. Percentages and iMFI were quantified and compared across conditions: control, HMBPP, HMBPP + apyrase, and PMA/ionomycin (left), or control, HMBPP, and PMA/ionomycin with or without BTN3A1 blocking antibody (0.1 µg/ml), and with or without IgG1 isotype control antibody (0.1 µg/ml) (right). Colored dots represent different donors. Statistical significance was calculated using ordinary one-way ANOVA. Asterisks indicate levels of statistical significance: *p < 0.05, **p < 0.01, ***p < 0.001; ns indicates a non-significant difference.

### Vγ9Vδ2 T Cell activation and IFNγ secretion require *Toxoplasma* invasion and BTN3A1 signaling

To determine whether parasite replication is necessary for IFNγ secretion, we infected PBMCs with the uracil-auxotrophic strain RHΔ*ompdc*Δ*up* under uracil-free conditions (Fig S2). These non-replicating parasites still triggered significant IFNγ secretion (Fig 2A). In contrast, pretreatment of the parasites with mycalolide B, which blocks actin-driven invasion (34), significantly reduced IFNγ release (Fig 2A), indicating that intracellular entry rather than replication *per se* is required.

**Fig 2.**
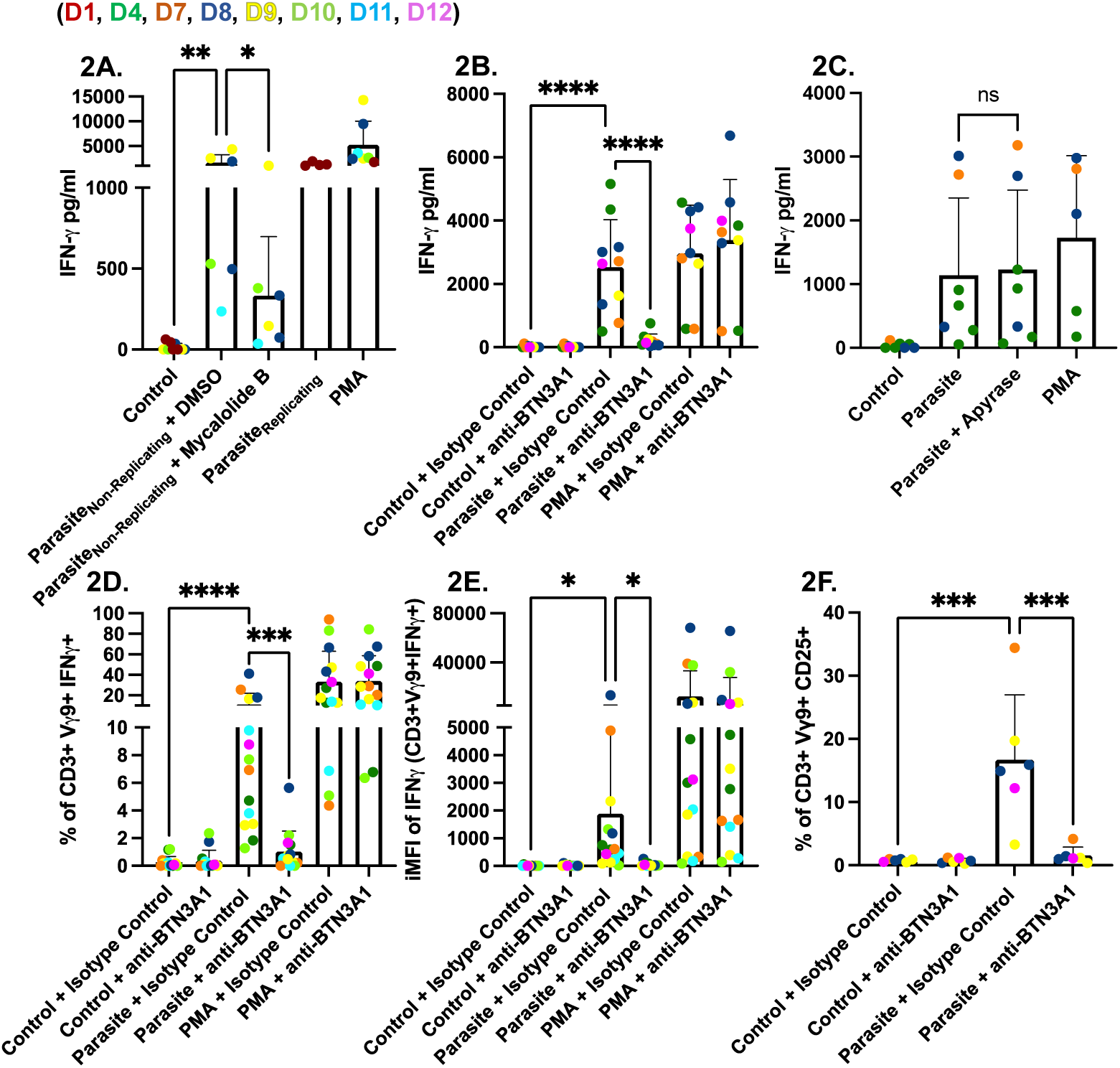
*Toxoplasma* stimulates Vγ9Vδ2 T cells to produce IFNγ via BTN3A1. **(A)** PBMCs (5×10^5^/well) were infected for 24 hours with non-replicating RHΔ*ompdc*Δ*up* parasites (non-replicating parasites) treated with or without Mycalolide B (10 µM) or with DMSO as a control, or infected with replicating parasites. PMA/ionomycin (1x) was used as a positive control, while unstimulated PBMCs were used as a negative control. IFNγ in culture supernatants was quantified by ELISA. **(B)** PBMCs were left untreated, infected for 24 hours with RHΔ*ompdc*Δ*up* parasites at an MOI of 0.6, or stimulated with PMA/ionomycin (1x). Cultures contained either BTN3A1-blocking antibody (0.1 µg/ml) or an IgG1 isotype control antibody (0.1 µg/ml), or no antibody. IFNγ release was measured as in panel A. **(C)** PBMCs were infected for 24 hours with RHΔ*ompdc*Δ*up* or RH wild-type parasites (MOI 0.6) with or without apyrase (200 IU/ml). PMA-stimulated and unstimulated cells served as controls. IFNγ concentrations (pg/ml) were determined by ELISA. **(D & E)** Intracellular cytokine staining of panel **D** shows the percentage of viable CD3^+^ Vγ9^+^ lymphocytes that are IFNγ-positive; panel **E** shows the (iMFI) of IFNγ within the Vγ9^+^ gate. **(F)** Activation marker analysis of the same samples: percentage of viable CD3^+^Vγ9^+^ lymphocytes expressing CD25. Data represent means + SD from multiple donors (individual data points shown). Statistical significance was calculated using one-way ANOVA followed by Tukey’s multiple comparison test. Asterisks indicate levels of statistical significance: *p < 0.05, **p < 0.01, ***p < 0.001, ****p < 0.0001; ns indicates a non-significant difference.

To limit PBMC death we utilized RHΔ*ompdc*Δ*up* in subsequent experiments. Infection with RHΔ*ompdc*Δ*up* induced high IFNγ levels, and these were significantly diminished by a BTN3A1-blocking antibody but not by an IgG1 isotype control (Fig 2B). Phosphoantigens from *Plasmodium falciparum* are liberated from infected red blood cells upon schizont rupture (17), raising the possibility that extracellular release might also occur here. We therefore added apyrase, which dephosphorylates HMBPP, during infections with RHΔ*ompdc*Δ*up* parasites. Apyrase had no appreciable effect on IFNγ production (Fig 2C), suggesting that the stimulatory phosphoantigen is retained within the infected cells and apyrase cannot access it. Flow cytometric analysis confirmed that the IFNγ detected by ELISA originates from Vγ9Vδ2 T cells. Infection increased both the percentage of CD3⁺ Vγ9⁺ lymphocytes producing IFNγ and their iMFI (Fig 2D,E). The same population up-regulated IL-2 alpha chain (CD25), an early activation marker (Fig 2F). In each case, BTN3A1 antibody blockade abolished the response, whereas the IgG1 isotype control antibody had no effect. Overall, these findings demonstrate that Vγ9Vδ2 T cells are the primary source of IFNγ in parasite-infected PBMCs cultures and their activation requires parasite invasion and signaling through the BTN3A1 receptor, but not intracellular parasite replication.

### Host, not parasite, isoprenoid biosynthesis drives Vγ9Vδ2 T cell IFNγ production during ***Toxoplasma* infection**

We sought to determine if parasite-derived or host-derived phosphoantigens were driving IFNγ induction. To determine if IFNγ production is driven by parasite-derived phosphoantigen we adapted an approach previously used for generating *Plasmodium falciparum* supernatants (17), and generated human foreskin fibroblasts (HFF) lysate and *Toxoplasma* RHΔ*ompdc*Δ*up* lysate that was filtered using a 3kDa filter to enrich for *Toxoplasma*-derived phosphoantigens. However, the 3kDa filtrate did not stimulate IFNγ secretion in PBMCs (Fig 3A). *Toxoplasma* lacks the mevalonate pathway and relies solely on the DOXP pathway within its apicoplast for isoprenoid precursor synthesis. Wild-type tachyzoites are intrinsically resistant to fosmidomycin, a competitive inhibitor of DOXP reductoisomerase (DOXPRI), because the drug does not cross the parasite plasma membrane and fails to accumulate at the enzymatic target (27). To bypass this permeability barrier, we used the GlpT strain, engineered to express the *E. coli* glycerol-3- phosphate/phosphate antiporter (GlpT) (27). Heterologous GlpT facilitates fosmidomycin uptake allowing DOXPRI inhibition, thereby limiting downstream production of HMBPP (Fig S3). Substitution of GlpT R45 to K and R269 to K (R45K-R269K) abolishes the activity of GlpT (27).

**Fig 3.**
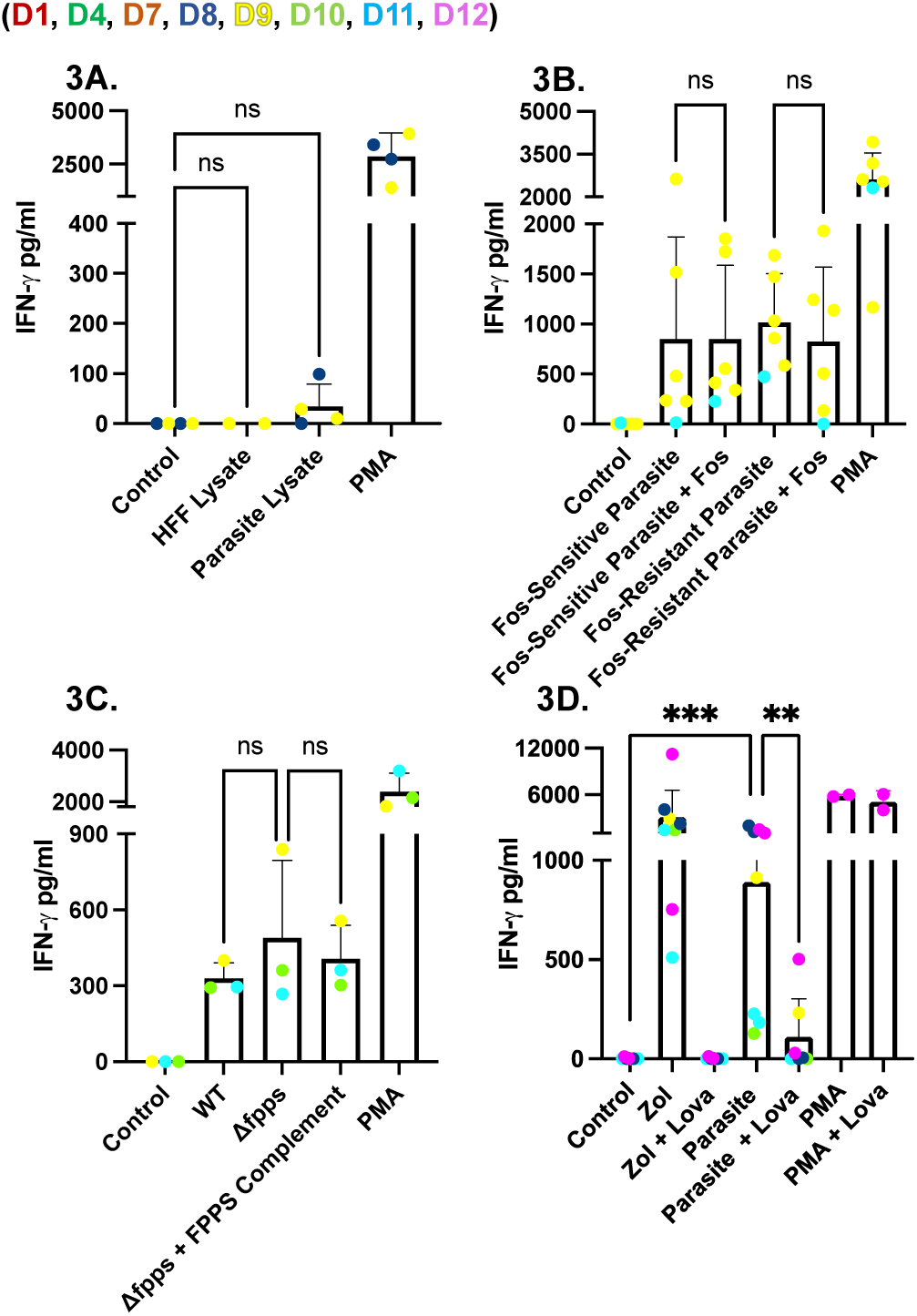
Cell-free parasite lysate, DOXP-pathway inhibition, and loss of farnesyl-pyrophosphate synthase do not reduce IFNγ release from *Toxoplasma*-infected PBMCs. (**A**) PBMCs (5×10^5^/well) were incubated for 24 hours with: HFF lysate or sonicated lysate from RHΔ*ompdc*Δ*up*-infected HFFs. The lysates were passed through a 3 kDa filter. PMA/ionomycin (1x) was used as a positive control, while unstimulated PBMCs were used as a negative control. **(B)** PBMCs were infected for 24 hours with GlpT (Fos-sensitive) or GlpT R45K-R269K (Fos- resistant)-expressing parasites with or without fosmidomycin (100 µM), an inhibitor of the DOXP pathway. **(C)** PBMCs were infected with RHΔ*hpt* (WT), RHΔ*hpt*Δ*fpps* (farnesyl-pyrophosphate-synthase knockout), or the corresponding complemented knockout strain for 24 hours. **(D)** PBMCs were pre-incubated with lovastatin (Lova) (25 µM) and/or zoledronate (Zol) (45 µM), washed three times, and then infected with RHΔ*ompdc*Δ*up* parasites for 24 hours. In all panels, IFNγ concentrations (pg/ml) in culture supernatants were measured by ELISA. Data represent means + SD from multiple donors (individual data points shown). Statistical significance was calculated using one-way ANOVA followed by Tukey’s multiple comparison test. Asterisks indicate levels of statistical significance: *p < 0.05, **p < 0.01, ***p < 0.001; ns indicates a non-significant difference.

Pre-treatment of GlpT (Fos-sensitive) and R45K-R269K (Fos-resistant) parasites with 100 µM fosmidomycin for 24 hours and subsequently using them to infect PBMCs led to no difference in IFNγ secretion (Fig 3B) even though plaque assays confirmed their susceptibility to fosmidomycin (Fig S4). Pre-treatment with 100 µM fosmidomycin for 48 hours and subsequently using them to infect PBMCs led to a reduction in IFNγ secretion; however, this drop was likely due to low parasite viability as demonstrated by plaque assays (data not shown). Deletion of farnesyl-pyrophosphate synthase (FPPS)(35), which lies immediately downstream of HMBPP, should allow HMBPP to accumulate while preventing flux into longer-chain isoprenoids (Fig S3). FPPS knockout parasites were validated by demonstrating their susceptibility to lovastatin (Fig S5), which inhibits host HMG-CoA reductase thereby preventing the FPPS knockout parasites from using host longer-chain isoprenoids, and then used to infect PBMCs. IFNγ secretion was not different between FPPS knockout and wild-type infection, arguing against an important role for parasite-derived HMBPP (Fig 3C).

Next, to assess the contribution of host isoprenoid biosynthesis, we treated infected PBMC cultures with lovastatin, a selective inhibitor of host 3-hydroxy-3-methylglutaryl-CoA reductase (HMG-CoA) (Fig S1). As a control we used zoledronate (Zol), an inhibitor of farnesyl pyrophosphate synthase (FPPS) (Fig S3), leading to accumulation of host IPP (36). Zoledronate led to robust IFNγ secretion while lovastatin significantly reduced IFNγ secretion. Similarly, lovastatin significantly reduced parasite-induced IFNγ secretion, consistent with a model in which BTN3A1-restricted Vγ9Vδ2 T cells respond primarily to host-generated IPP rather than to parasite- derived HMBPP (Fig 3D). Overall, these results indicate that IFNγ production requires an active intracellular parasite infection but depends more critically on the host mevalonate pathway than on the parasite DOXP pathway.

### Vγ9Vδ2 T Cells mediate BTN3A1-dependent activation of NK Cells via TNFα to promote IFNγ responses to *Toxoplasma*

To determine if IFNγ is produced by cell populations other than Vγ9Vδ2 T cells, PBMCs were gated on CD3^+^Vγ9^-^-lymphocytes (mainly conventional CD4 and CD8 T cells) and on CD3^-^ - lymphocytes (predominantly NK cells) (13) within our population. We saw no differences in either the percentage or iMFI of CD3^+^Vγ9^-^IFNγ^+^ cells between our control and infected PBMCs (Fig 4A and 4B). In contrast, we saw a significant increase in the percentage (Fig 4C) and iMFI (Fig 4D) of CD3^-^IFNγ^+^ cells in parasite infected PBMCs; this increase was significantly reduced by an anti- BTN3A1 blocking antibody but not by IgG1 isotype control. To confirm that CD3^-^IFNγ^+^ cells were NK cells, we used the CD56 marker and again observed a significant increase in both the percentage (Fig 4E) and iMFI (Fig 4F) of CD3^-^CD56^+^IFNγ^+^ cells in infected PBMCs. This increase was reduced when BTN3A1 blocking antibody was added to the cultures. These results indicate that NK cell IFNγ production during *Toxoplasma* infection is dependent on BTN3A1 signaling, suggesting that NK cells act as secondary responders following initial Vγ9Vδ2 T cell activation.

**Fig 4.**
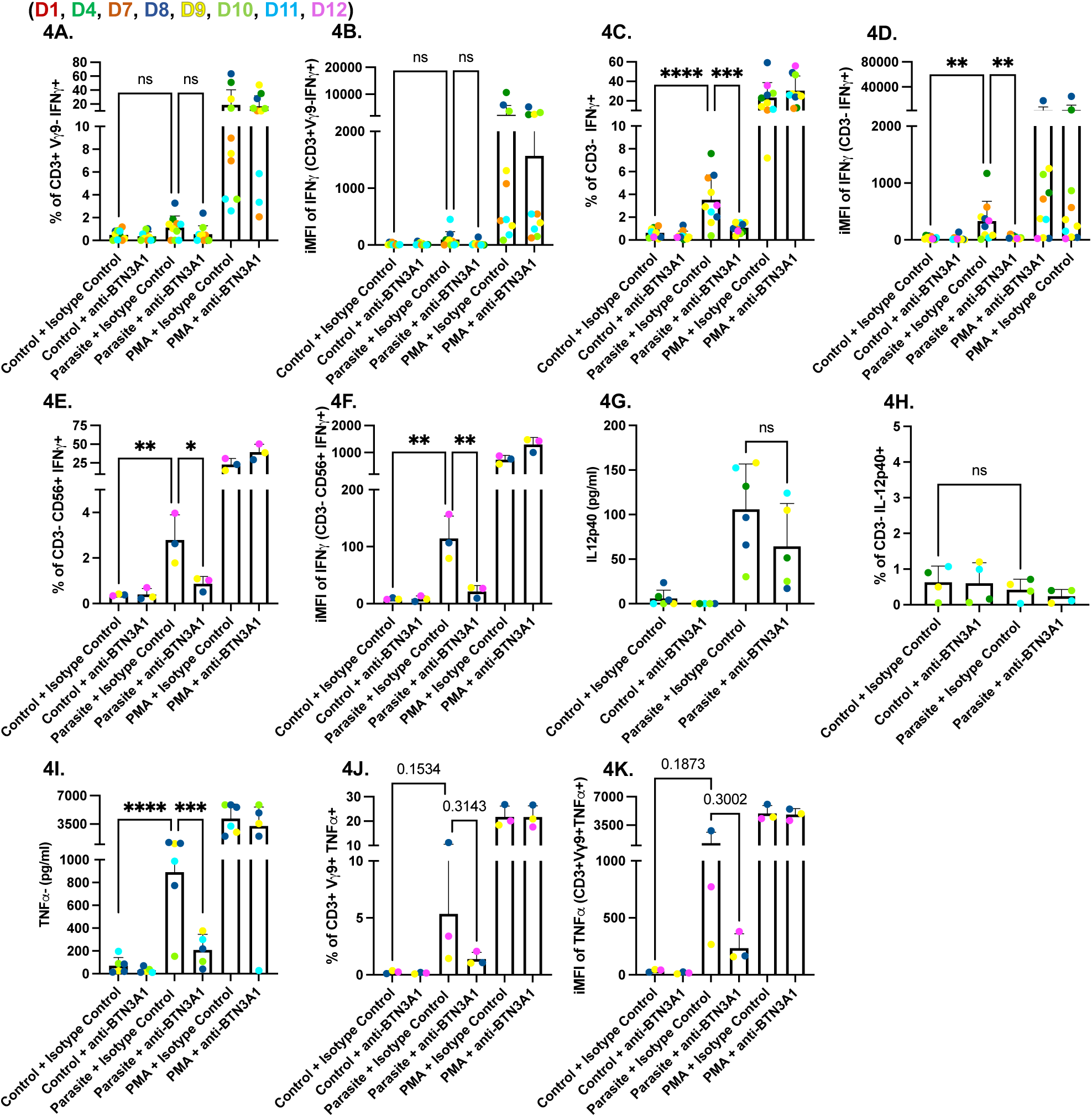
BTN3A1 signaling is required for IFNγ production by NK cells in parasite-infected PBMCs. PBMCs (5×10^5^ cells/well) were left untreated, infected for 24 hours with the RHΔ*ompdc*Δ*up* strain (parasite) at an MOI of 0.6, or stimulated with PMA/ionomycin (1x). Each condition was cultured with BTN3A1-blocking antibody (0.1 µg/ml), or with IgG1 isotype control antibody (0.1 µg/ml). PBMCs were gated on live, single lymphocytes. **(A)** Percentage of CD3^+^/Vγ9^-^/IFNγ^+^ cells **(B)** iMFI of IFNγ in the CD3^+^/Vγ9^-^ subset **(C)** Percentage of CD3^-^/IFNγ^+^ cells. **(D)** iMFI of IFNγ in CD3^-^ cells. **(E)** Percentage of CD3^-^/CD56^+^/IFNγ^+^ cells **(F)** iMFI of IFNγ in CD3^-^/CD56^+^/IFNγ^+^cells **(G)** IL-12p40 and (**H**) TNFα concentrations (pg/ml) in culture supernatants measured by ELISA. Data represent means + SD from multiple donors (individual data points shown). Statistical significance was calculated using one-way ANOVA followed by Tukey’s multiple comparison test. Asterisks indicate levels of statistical significance: *p < 0.05, **p < 0.01, ***p < 0.001, ****p < 0.0001; ns indicates a non-significant difference.

This led us to hypothesize that Vγ9Vδ2 T cells drive IFNγ secretion from NK cells through a BTN3A1-dependent release of TNFα that acts in concert with IL-12 secreted by infected monocytes (37). Consistent with this model, parasite infection increased IL-12 in culture supernatants, and anti-BTN3A1 had no effect on this increase (Fig 4G). However, we saw no differences in IL-12 secretion in CD3^-^ cells (Fig 4H).

In contrast, TNFα secretion was robust in *Toxoplasma*-infected PBMCs but was significantly reduced when BTN3A1 signaling was blocked (Fig 4I). Intracellular cytokine staining confirmed that Vγ9Vδ2 T cells produced most of this cytokine: parasite infection increased both the frequency of TNFα-positive Vγ9Vδ2 T cells (Fig 4J) and their iMFI (Fig 4K), whereas BTN3A1 inhibition returned both measures to baseline. Together, these data indicate that while Vγ9Vδ2 T cells are the primary BTN3A1-dependent responders to *Toxoplasma* infection, the TNFα they secrete may reinforce IL-12-driven NK-cell activation, leading to a secondary wave of IFNγ production.

### *Toxoplasma* promotes BTN3A1-dependent expansion of human Vγ9Vδ2 T cells

Previous studies have demonstrated an expansion of Vγ9Vδ2 T cells in infants of mothers infected with *Toxoplasma* (29). Consistent with previous data, both HMBPP (Fig 1A) and *Toxoplasma* (Fig 2A) were capable of stimulating IFNγ production from Vγ9Vδ2 T cells, which was completely abrogated by a BTN3A1-blocking antibody. We next investigated whether *Toxoplasma*, in combination with IL-2, could drive the clonal expansion of Vγ9Vδ2 T cells and whether this expansion can be inhibited with BTN3A1 blocking antibodies.

Human PBMCs were cultured with IL-2 and stimulated with HMBPP or infected with a low MOI of RHΔ*ompdc*Δ*up* to minimize cell death (Fig 5A). After two stimulation cycles, the frequency of Vγ9+ cells within the CD3+ compartment increased approximately two-fold across all donors examined (Fig 5C). This expansion was smaller than the six-fold increase observed with HMBPP (Fig 5B). Inclusion of the BTN3A1 blocking antibody throughout the culture period reduced Vγ9Vδ2 T cell frequencies to baseline, confirming that expansion is strictly BTN3A1-dependent (Fig 5C). Although the frequency of CD3^+^Vγ9^+^ cells varied among donors, expansion was evident in response to parasites compared to conditions of BTN3A1 antibody or control. Overall, our data demonstrate that human Vγ9^+^ T cells expand in response to *Toxoplasma* and IL-2 and that this process requires the BTN3A1 receptor.

**Fig 5.**
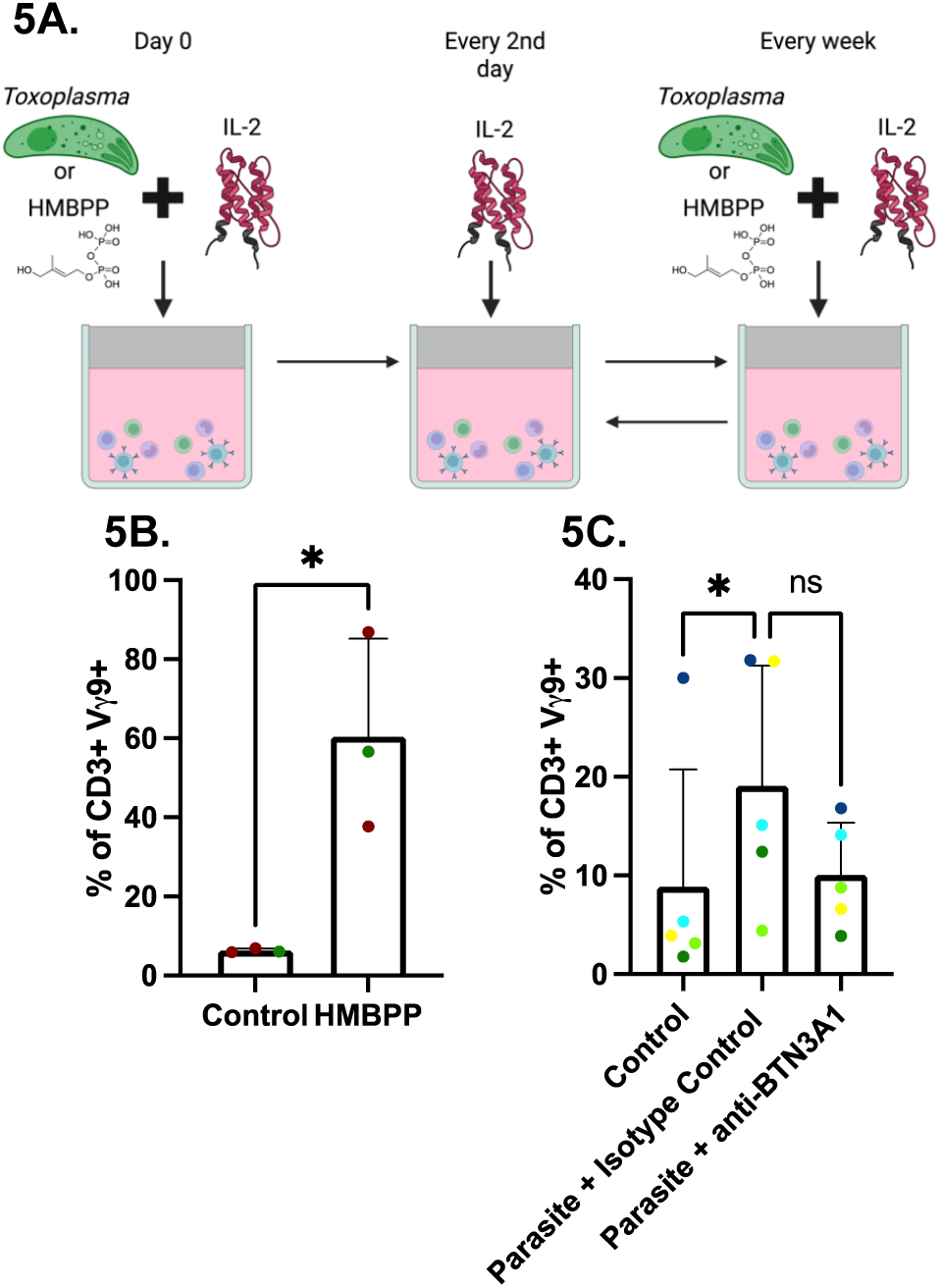
BTN3A1-dependent expansion of Vγ9Vδ2 T cells following exposure to *Toxoplasma* or HMBPP. **(A)** Experimental timeline. PBMCs (1×10^6^) were infected with RHΔ*ompdc*Δ*up* parasites at an MOI of 0.1 in uracil-free medium, supplemented with IL-2 (50 IU/ml) and either BTN3A1 blocking antibody (0.1 µg/ml) or IgG1 isotype control. IL-2 was replenished every other day until day seven, when cells were washed and stimulated or reinfected under identical conditions. After day 14, only IL-2 was added and at day 21 cells were analyzed by flow cytometry. Figure was created in BioRender. Rodriguez, F. (2025) https://BioRender.com/661lzno. Percentage of lymphocytes/single cells/alive/CD3+ in Vγ9^+^ cells in HMBPP- (**B**) or parasite- stimulated cells (**C**). Data represent means + SD from multiple donors (individual data points shown). Statistical significance was calculated using t-test (**B**) or non-parametric test followed by Dunn’s multiple comparison test (**C**). Asterisks indicate levels of statistical significance: *p < 0.05; ns indicates a non-significant difference.

## Discussion

While previous studies have indirectly suggested a role for Vγ9Vδ2 T cells during *Toxoplasma* infection, our findings provide direct evidence that Vγ9Vδ2 T cells are the dominant early source of IFNγ in parasite-infected human PBMCs. Activation of these cells requires BTN3A1 and depends largely on host-derived, mevalonate pathway phosphoantigens rather than on parasite-encoded metabolites. Furthermore, we demonstrate that Vγ9Vδ2 T cells enhance NK cell IFNγ production in a BTN3A1-dependent manner, likely through a synergistic effect of TNFα from Vγ9Vδ2 T cells and IL-12 and IL-1β secreted from infected cells.

Despite effective inhibition of exogenous HMBPP by apyrase, this enzyme failed to suppress IFNγ secretion induced by *Toxoplasma* infection. Similarly, shrimp alkaline phosphatase failed to prevent *Toxoplasma-*induced IFNγ secretion (data not shown). These enzymes are membrane-impermeant and therefore cannot access intracellular phosphoantigens, supporting the idea that the relevant ligand accumulates inside infected host cells. Contrary to *P. falciparum* and *L. monocytogenes* infection, where lysates effectively stimulate Vγ9Vδ2 T cells (17,38), *Toxoplasma* lysate failed to stimulate. A possible explanation might be that *Toxoplasma* NTPases with apyrase-like activity degrade parasite-derived phosphoantigens (39), which could lower the amount of HMBPP that reaches the host cytosol, reducing binding to BTN3A1, and consequently limiting Vγ9Vδ2 T cell activation.

Chemical and genetic probing confirmed that host, not parasite, isoprenoid biosynthesis is critical for Vγ9Vδ2 T cell activation. Lovastatin, an HMG-CoA-reductase inhibitor, blocked IFNγ production, whereas zoledronate, which blocks farnesyl-pyrophosphate synthase and causes IPP build-up, enhanced it. Transcriptomic datasets from *Toxoplasma*-infected HFFs and PBMCs show coordinated upregulation of mevalonate enzymes (3-hydroxy-3-methylglutaryl-CoA synthase (HMGCS1), mevalonate kinase (MVK), 3-hydroxy-3-methylglutaryl-CoA reductase (HMGCR), MVD, and isopentenyl-diphosphate delta isomerase (IDI1)(Table S1)(40,41), suggesting that *Toxoplasma* actively boosts IPP synthesis to levels sensed by BTN3A1-expressing host cells. Additionally, geranylgeranyl diphosphate synthase is downregulated, potentially limiting IPP consumption and thereby contributing to its accumulation (Table S1)(40). *E. coli* and *S. aureus* infections synergized with zoledronate to enhance TNFα secretion from Vγ9Vδ2 T cells, while addition of mevastatin abrogated this response entirely (42). Host PP2A phosphatase-dependent dephosphorylation of HMGCR leading to its activation has been implicated in Vγ9Vδ2 T cell activation (42). However, PP2A subunit expression did not change significantly in PBMCs (40)(Table S1), indicating *Toxoplasma* may use alternative mechanisms. A possible mechanism could be through the effector GRA16, which is secreted beyond the vacuole, and has been shown to increase expression of PP2A-B55 via NF-kB, leading to cell cycle arrest and apoptosis in lung carcinoma cells (43).

HMBPP-deficient *L. monocytogenes* mutants can still activate Vγ9Vδ2 T cells when infecting human monocyte-derived dendritic cells, while lysates alone failed to activate these T cells (38). Notably, infection upregulated host mevalonate/cholesterol pathway genes like HMGCR, reinforcing the concept that host IPP is a key danger signal. Transporters such as ABCA1 and APOA1 have been proposed to export IPP (44) and ABCA1, but not APOA1 expression, is reduced in human dendritic cells when mevastatin is used during an infection with HMBPP- deficient *L. monocytogenes* mutants (38). However, ABCA1 expression was not significantly modulated, while APOA1 expression was too low to be detected (Table S1)(40).

BTN3A1 blockade eliminated both Vγ9Vδ2 T cell-derived IFNγ and the IFNγ from NK cells yet had no effect on IL-12 release. These observations support a model wherein Vγ9Vδ2 T cells license NK cells to secrete IFNγ during *Toxoplasma* infection. We proposed a cytokine-mediated response where IL-12 from infected cells and TNFα from Vγ9Vδ2 T cells substantially enhance NK cell-derived IFNγ secretion (Fig 6). This model aligns with previous studies indicating a synergistic role of TNFα and IL-12 in amplifying NK cell activity (37,45). Parasite effectors GRA15 and GRA24 are known to activate host NF-kB and p38 MAPK, respectively, promoting IL-12 and IL-1β synthesis by monocytes and dendritic cells (46–48). IL-1β and IL-12 have been demonstrated to amplify IFNγ in NK cells during *Toxoplasma* infection (49). Future work should test whether blocking TNFα diminishes NK cell licensing and compromises early parasite control.

**Fig 6.**
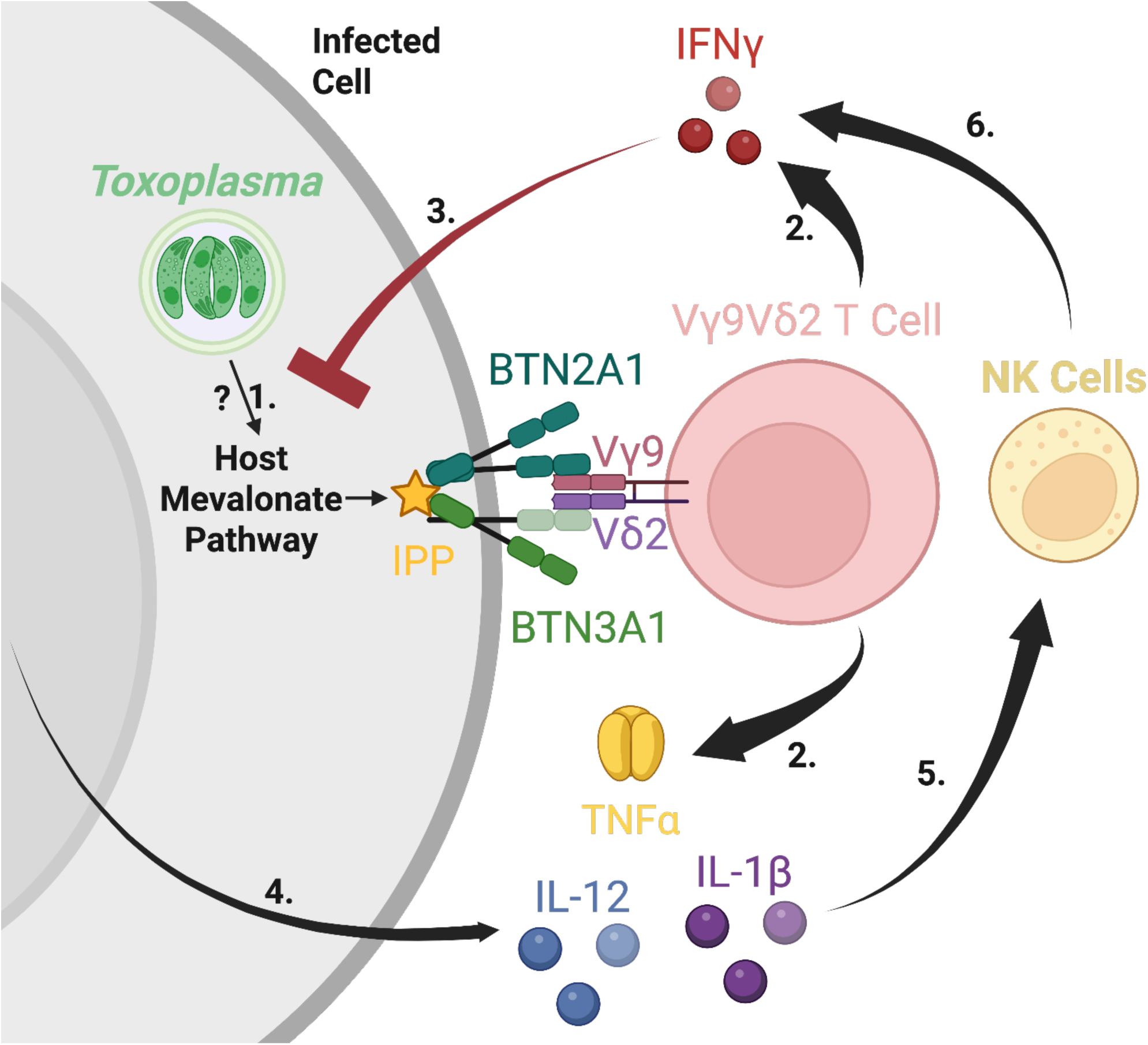
Accumulation of IPP through the mevalonate pathway activates BTN3A1 and coordinates Vγ9Vδ2 T-cell and NK-cell interferon gamma responses during *Toxoplasma* infection. 1. *Toxoplasma* infection upregulates host-cell mevalonate-pathway enzymes, leading to the accumulation of host cell isopentenyl pyrophosphate (IPP) promoting activation of BTN3A1/2A1 dual receptor. **2.** Vγ9Vδ2 T cells recognize IPP-activated BTN3A1/2A1 leading to their activation and secretion of IFNγ and TNFα. **3.** IFNγ stimulates infected cells by binding to the IFNγ receptor leading to upregulation of multiple interferon-stimulated genes that encode anti-parasitic effectors. **4.** IL-12 and IL-1β get secreted by infected cells **5.** IL-12 and IL-1β synergize with TNFα from Vγ9Vδ2 T cells to activate NK cells. **6.** Activated NK cells secrete additional IFNγ thereby amplifying the anti-parasite response. Model was created in BioRender. Rodriguez, F. (2025) https://BioRender.com/8i8ocqt

Overall, our findings challenge the notion that Vγ9Vδ2 T cell activation is solely due to parasite-derived HMBPP. Instead, the parasite manipulates host lipid metabolism, which generates sufficient IPP for BTN3A1 engagement. Vγ9Vδ2 T cells then act as sentinels, supplying rapid IFNγ and TNFα that augment NK cell IFNγ secretion. This layered response may compensate for the absence of TLR11/12 and IRG pathways in humans. Defining the parasite effectors and host sensors that modulate mevalonate flux will be essential for understanding, and potentially manipulating, early immunity to *Toxoplasma* and other apicomplexans.

## Acknowledgements

RHΔ*ompdc*Δ*up* was obtained from Dr. David J. Bzik (34). GlpT and R45K-R269K parasite strains were obtained from Dr. Boris Striepen (27). RHΔ*ku80*Δ*fpps*, RHΔ*ku80*Δ*fpps +* FPPS, and WT parasite strains were obtained from Dr. Silvia Moreno (35). This project was supported by the UC Davis Flow Cytometry Shared Resource Laboratory with funding from NCI P30 CA093373 and James B. Pendleton Charitable Trust: BD “LSRII” Cytometer (Davis) with technical assistance from Ms. Bridget Mclaughlin. Research reported in this publication was supported by the National Institute of Allergy and Infectious Diseases of the National Institutes of Health under Award Number R01AI173803. The content is solely the responsibility of the authors and does not necessarily represent the official views of the National Institutes of Health. Felipe Rodriguez received fellowships from the Graduate Student Support Program (GSSP) supported by the Hart endowed fellowship and Vivian and Dorothea Hagaman fellowship. Additionally, he received a UC Davis internal fellowship supported by Stephen F. and Bettina A. Sims Immunology fellowship.

## Ethics Statement

The present study involved the utilization of fresh blood samples from healthy anonymous donors, which were provided by Vitalant, a not-for-profit community blood center. Vitalant received IRB approval from the Western Institutional Review Board under the protocol BSI-18-001. The supplier holds licenses from the U.S. Food and Drug Administration (FDA) and complies with all Clinical Laboratory Improvement Amendments (CLIA) regulatory requirements. Additionally, Vitalant is accredited by the AABB (formerly known as the American Association of Blood Banks). Throughout the research process, strict adherence to ethical guidelines was maintained to ensure the confidentiality and anonymity of the donors.

## Material and Methods

### Antibodies and reagents

The BTN3A1 blocking antibody (clone 103.2) was from Creative Biolabs, and an IgG1 isotype control antibody was from Sigma. (E)-1-Hydroxy-2-methyl-2-butenyl 4-pyrophosphate lithium salt (HMBPP)(95098-1MG) and potato apyrase (A6535-100UN) was from Sigma, while isopentenyl diphosphate (IPP)(I-0050) was from Echelon Biosciences. All fluorescent and intracellular staining antibodies were from Biolegend: Brilliant Violet 711 anti-human CD3, Brilliant Violet 421 anti-human CD25, PE/Cyanine5 anti-human CD56, PE anti-human TCR Vγ9, APC anti-human IFNγ, FITC anti-human IL-12p40, BV650 anti-human TNFα, and Human TruStain FcX. Live Dead dye: Zombie NIR Live/Dead and PMA/ionomycin (500x) was from Biolegend. Staining protocols were as directed by the manufacturer (Biolegend). Ficoll-Paque Premium (17-5442-02) was from GE Healthcare. Mycalolide B (BML-T123) was from Enzo. Zoledronate disodium salt (ab143738) was from Abcam. Costar 96 well plates (3361) for ELISA were from Corning. Human blood was collected by Vitalant. Matched-pair human ELISA kits IFNγ (CHC1233), TNFα (CH1753), or IL-12p40 (CHC1563) were from Thermo Fisher.

### Culture of cells and parasites

Human foreskin fibroblasts (HFFs) were cultured at 37°C at 5% CO_2_ in Dulbecco’s Modified Eagle Medium (DMEM; Invitrogen) supplemented with 10% heat-inactivated fetal bovine serum (HIFBS), 2 mM of L-glutamine, 50 µg/ml penicillin/streptomycin, and 20 µg/ml gentamicin (HFF medium). Parasites were cultured similarly in DMEM supplemented with 1% HIFBS, 50 µg/ml penicillin/streptomycin, and 2 mM of L-glutamine (*Toxoplasma* medium). RHΔ*ompdc*Δ*up* (uracil auxotroph strain) was grown in *Toxoplasma* medium containing 250 µM uracil (Uracil medium). Human PBMCs were cultured in RPMI-1640 medium supplemented with 25 mM HEPES, 2 mM L-glutamine, 20 µg/ml gentamicin, 50 µg/ml penicillin/streptomycin, and 10% HIFBS (PBMC media).

### PBMC isolation

Donor blood was added into 50 ml conical tubes and centrifuged for 20 mins at 500 x g at room temperature (RT) (no acceleration or brake). A serum aliquot was taken to test for *Toxoplasma* specific antibodies. The buffy coat was then transferred into a 15 ml conical tube containing PBS and mixed thoroughly before carefully layering over Ficoll-Paque (1:1, Histopaque-1077) and centrifuged at 680 x g for 30 mins at RT (no acceleration or brake). The PBMC interface was collected, washed in ice cold PBS, and spun at 200 x g for 10 mins at 4°C. The cell pellet was resuspended in PBMC media. Cell viability, determined by trypan blue exclusion, confirmed that over 95% of the cells were viable. PBMCs were cryopreserved in 90% FBS/10% dimethyl sulfoxide (DMSO) for later use.

### Serum test for *Toxoplasma* specific antibodies

High-binding 96-well plates were coated overnight at 4°C with lysate (0.1 μg/well) in 0.1 M bicarbonate buffer (pH 9.6). Plates were washed with PBS-T (0.05% Tween-20) and blocked for ≥1 h at room temperature with 2% casein in PBS. Human sera were diluted (1:40, 1:80, 1:160) in 1% casein, added in duplicate (50 μl/well), and incubated for 1–2 hours at room temperature. After washing, wells were incubated with HRP-conjugated mouse anti-human IgG (BD Pharmingen, 1:1000) for 1 hour in the dark. ABTS substrate (KPL) was added, and plates were read at 405 nm at 20 and 40 min (lysates). The optical density (OD) obtained with *Toxoplasma* lysate was divided by the mean OD values of *Sarcocystis neurona* and *Neospora hughesi*. A positive signal was defined as OD > 1.4.

### Determination of intracellular parasite growth

Parasite per vacuole counts were performed using 24-well plates with confluent HFF monolayers and infected with GlpT and R45K-R269K parasites with/out 100 µM fosmidomycin incubated for an hour prior to infection while RHΔ*ompdc*Δ*up* were incubated with 250 µM uracil. Parasites were fixed in 3% paraformaldehyde and stained with SAG2A and DAPI. Parasites per vacuole were quantified at 24 and 48 hours after infection and at least 100 vacuoles per strain and per condition were counted using microscopy.

### Expansion of Vγ9Vδ2 T cells

Frozen PBMCs were thawed in a 37°C water bath, subsequently spun, washed, and counted/checked for viability. The thawed PBMCs were then stimulated with HMBPP (312 nM) and IL-2 (200 U/ml) to expand Vγ9Vδ2 T cells. In parallel, a uracil-auxotroph parasite strain was used to expand Vγ9Vδ2 T cells in medium without uracil. Cells were seeded in a 24-well flat bottom plate and cultured with IL-2 (50 IU/mL) replenished every second or third day. Cultures were split when culture medium acidification (yellowing) was observed, usually between day six and day eight, by replacing the medium with fresh medium with IL-2, followed by re-seeding the cells. After the second week of culture, only IL-2 was added to minimize IFNγ production. The expansion of the Vγ9Vδ2 T-cells was carried out for approximately 21 days.

### Intracellular cytokine staining

PBMCs were processed using FACS buffer (PBS, pH 7.4, with 2%HIFBS). Cells were resuspended at a concentration of 5 x 10^6^ cells/ml in PBMC media and plated in a 96-well round-bottomed tissue culture plate at 5 x 10^5^ cells per well (100 μl of cell suspension) and incubated at 37°C at 5% CO_2_. Single-stain and fluorescence-minus-one controls were included. Cells were stimulated with HMBPP (312 nM) or infected using RHΔ*ompdc*Δ*up* (0.6 MOI) without uracil, or with BTN3A1 blocking antibody or IgG1 isotype control antibody (0.1 µg/ml). All wells after infection or stimulation had a final volume of 200 µl. After incubating for 16 hours, PMA/ionomycin (1x) was added to positive control wells and GolgiStop to all wells (0.067%/1:1500) (BD Biosciences) for the last 8 hours. After 24 hours, the plate was then processed for staining. PBMCs were washed with 1x PBS containing 1x EDTA and stained with Zombie NIR for live/dead discrimination. The plate was then placed in ice and washed with FACS buffer. Human TruStain FcX was used to block as directed by manufacturer (Biolegend). For surface staining, a cocktail of surface-marker antibodies was used (Antibodies and reagents section). For intracellular staining, the cells were fixed and permeabilized using Cyto-Fast™ Fix/Perm Buffer Set (Biolegend), APC anti-human IFNγ, FITC anti-human IL-12p40, or BV650 anti-human TNFα was used as the intracellular marker. After staining, the cells were washed and resuspended in the FACS buffer for analysis on the flow cytometer.

### *In vitro* cytokine ELISA

PBMCs were resuspended at a concentration of 5 x 10^6^ cells/ml in PBMC media and plated in a 96-well round-bottomed tissue culture plate at 5 x 10^5^ cells per well (100 μl of cell suspension) and incubated at 37°C at 5% CO_2_. PBMCs were stimulated with HMBPP (312 nM) or infected using RHΔ*ompdc*Δ*up* (0.6 MOI) without uracil, with or without apyrase (200 IU/ml), or with BTN3A1 blocking antibody or IgG1 isotype control antibody (0.1 µg/ml). For zoledronate and lovastatin experiments, PBMCs were treated with lovastatin (25 µM) for 1 hour, prior to stimulating with zoledronate (45 µM), RHΔ*ompdc*Δ*up* (0.6 MOI), or PMA/ionomycin (1x) for 24 hours. Supernatants were collected after 24 hours and used to determine IFNγ, TNFα, or IL-12p40 levels. All the cytokine levels were measured using commercially available matched-pair human ELISA kits (Thermofisher; CHC1233 (IFNγ), CH1753 (TNFα), CHC1563 (IL-12p40), following the manufacturer’s instructions.

### Glycerol 3-transporter confirmation in *Toxoplasma*

GlpT and R45K-R269K parasite strains were confirmed to be mycoplasma-free using GPO-3 and MGSO primers (Table S2). To confirm type I strain, GRA6 FW and RV was used with restriction enzyme MseI, while glycerol 3-transporter was confirmed with GlpT FW1 and GlpT RV1 (Table S2). Glycerol-3-phosphate (G3P)-transporter was tagged by TY (24), immunofluorescence assay was performed to confirm localization of the G3P transporter in GlpT and R45K-R269K parasites, RHΔ*hpt* was used as a control. Staining was done using mouse-anti-Ty, rabbit-anti-SAG2A (parasite surface marker), and DAPI. Glycerol 3-transporter amplicon was sequenced using GlpT FW1 and GlpT FW2 by Quintara Bio. Amino acid changes were confirmed in the glycerol 3-transporter sequence (arginine to lysine in positions 45 and 269 in the R45K-R269K parasite strain).

### Statistics

All statistical analyses were performed using Prism (GraphPad) version 10.3.1. Data are presented as mean ± standard deviation (SD). Statistical significance was determined using one-way ANOVA with Tukey’s post hoc test for comparisons involving more than two groups, and unpaired *t*-tests or non-parametric tests with Dunn’s multiple comparisons for two-group analyses, as specified in figure legends. Significance was defined as *p* < 0.05 and denoted by asterisks in all figures.

---

**Fig S1.**
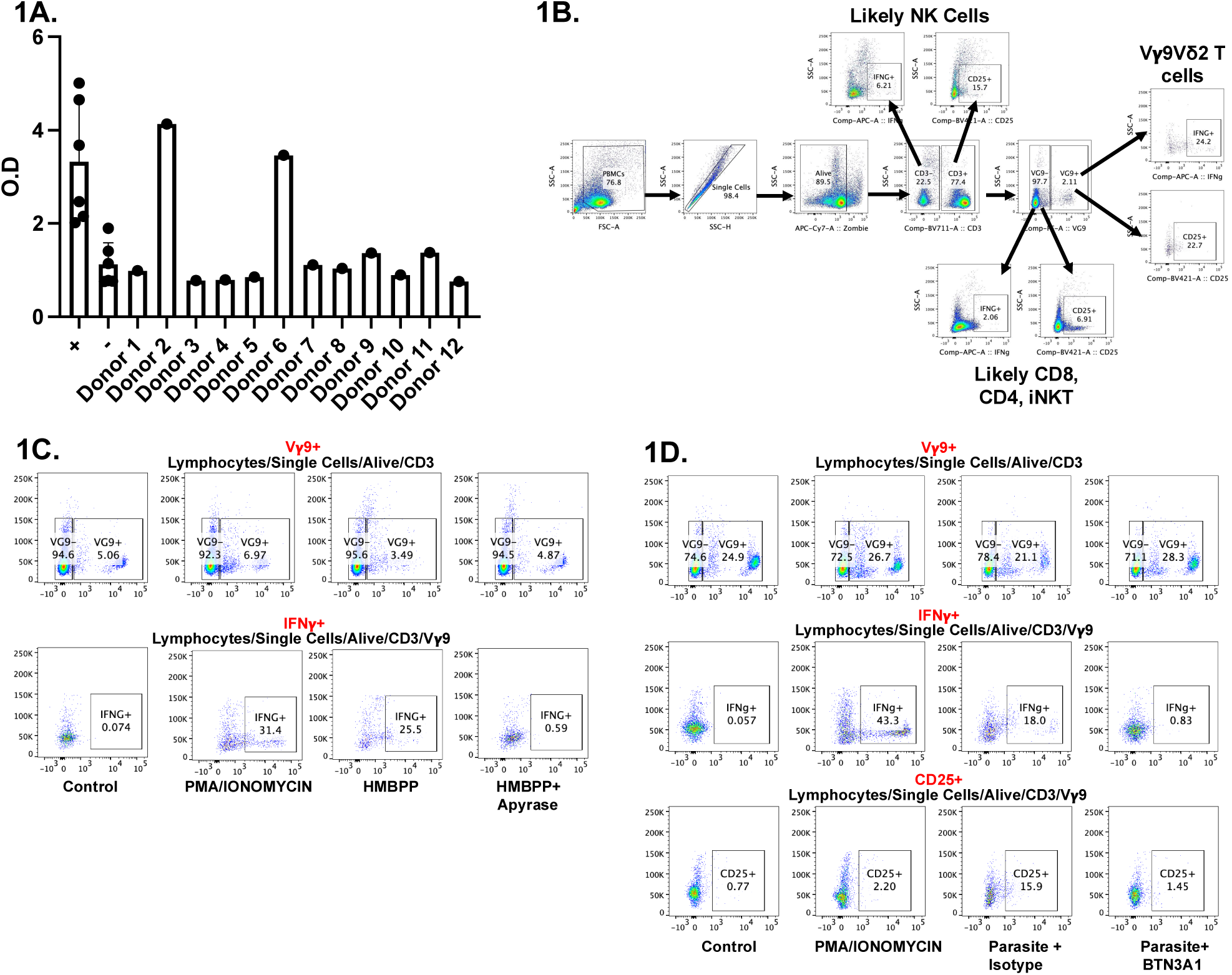
Serological screening of blood donors and flow-cytometric gating strategy for PBMCs. **(A)** Donor sera were tested for *Toxoplasma*-specific antibodies by ELISA using sonicated *Toxoplasma* lysate as the test antigen and lysates of *Sarcocystis neurona* and *Neospora hughesi* as negative controls. The optical density (OD) obtained with *Toxoplasma* lysate was divided by the mean OD of the two control lysates to generate a reactive ratio for each donor. Donors 2 and 6 were not used as they were considered *Toxoplasma* positive (**B)** Flow-cytometry gating strategy used for identifying cell populations. PBMC events were sequentially gated in FlowJo as follows: 1. Lymphocytes (SSC-A vs. FSC-A), 2. Single cells (SSC-A vs. SSC-H), 3. Viable cells (SSC-A vs. APC-Cy7::Zombie NIR), 4. CD3^+-^ (SSC-A vs. BV711::CD3), 5. Vγ9^+-^ (SSC-A vs. PE::Vγ9), 6. IFNγ^+^ (SSC-A vs. APC::IFNγ) or CD25^+^ (SSC-A vs. BV421::CD25). (**C-D**) Dot plots show the percentage of Vγ9^+^ cells within the live CD3^+^ gate and the percentage of IFNγ^+^ cells within the live CD3^+^/Vγ9^+^ gate under the following conditions: unstimulated, (**C**) HMBPP (312 nM), HMBPP + apyrase (200 IU/ml), and PMA/ionomycin (1x). (**D)** Infection for 24 h with RHΔ*ompdc*Δ*up* parasites (MOI 0.6) with or without BTN3A1 blocking antibody or IgG1 isotype control antibody (0.1 µg/ml).

**Fig S2.**
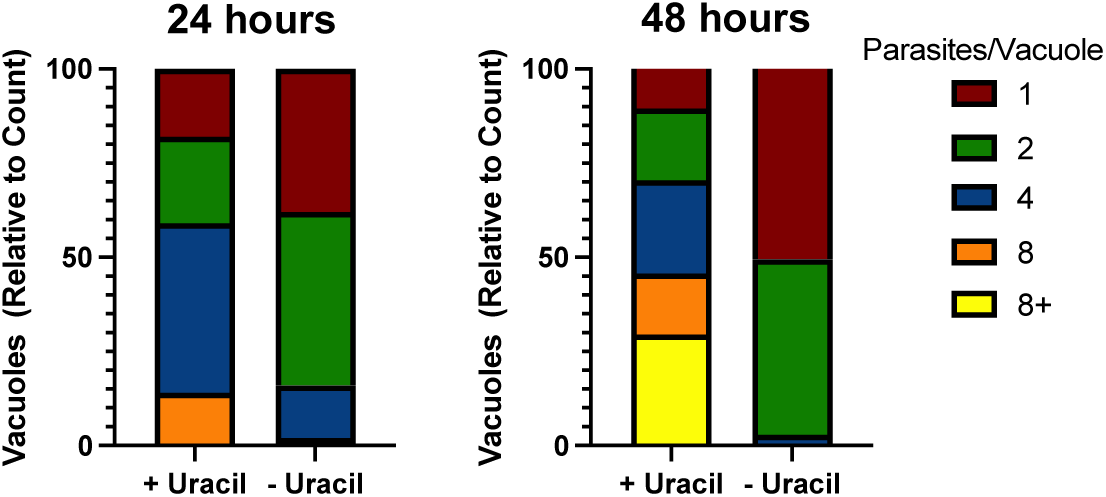
RHΔ*ompdc*Δ*up* parasites have impaired growth without uracil. Parasite replication was assessed by counting parasites per vacuole in RHΔ*ompdc*Δ*up*-infected HFF monolayers cultured with or without 250 µM uracil. Parasite counts were performed at 24 hours (left) or 48 hours (right) post-infection. Samples were fixed with 3% paraformaldehyde and stained with anti-SAG2A (parasite surface marker) and DAPI. Parasites per vacuole were quantified by fluorescence microscopy.

**Fig S3.**
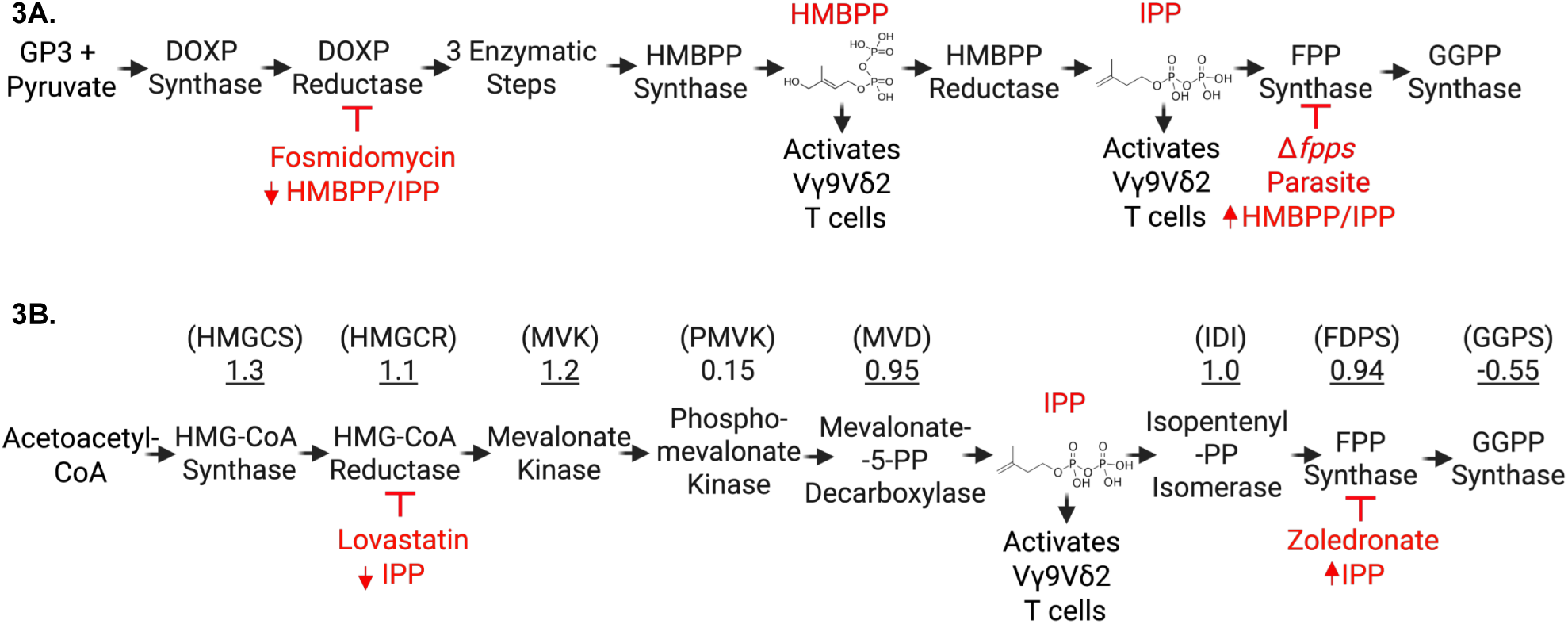
Comparative overview of the apicoplast 1-deoxy-D-xylulose-5-phosphate (DOXP) pathway and the host cytosolic mevalonate (MEV) pathway. **A)** The non-mevalonate DOXP pathway operates in apicoplast-containing protozoa such as *Toxoplasma* and *Plasmodium spp.*, most bacteria, and in chloroplasts. Sequential reactions convert 1-deoxy-D-xylulose-5-phosphate to the phosphoantigens hydroxymethyl-butenyl pyrophosphate (HMBPP) and isopentenyl pyrophosphate (IPP). Fosmidomycin blocks DOXP reductoisomerase, halting production of both intermediates. Deletion of farnesyl-pyrophosphate synthase (Δ*fpps*) truncates the pathway downstream of IPP/HMBPP, leading to their intracellular accumulation. Created in BioRender. Rodriguez, F. (2025) https://BioRender.com/yba9idv **B)** Eukaryotes synthesize IPP from acetyl-CoA via the mevalonate route. The rate-limiting step, catalysed by HMG-CoA reductase (HMGCR), is inhibited by lovastatin, preventing IPP formation. Farnesyl-pyrophosphate synthase (FPPS) condenses IPP with dimethylallyl-PP; zoledronate inhibits FPPS, causing upstream IPP build-up. Gene IDs above each enzyme are based on RNA-seq data from the GEO dataset GSE119835 (Table S1). Numbers indicate the log2Fold change expression and underlined values denote statistical significance (adjusted p < 0.05, Benjamini-Hochberg FDR) (Table S1). Figure S3 was created in BioRender. Rodriguez, F. (2025) https://BioRender.com/yba9idv

**Fig S4.**
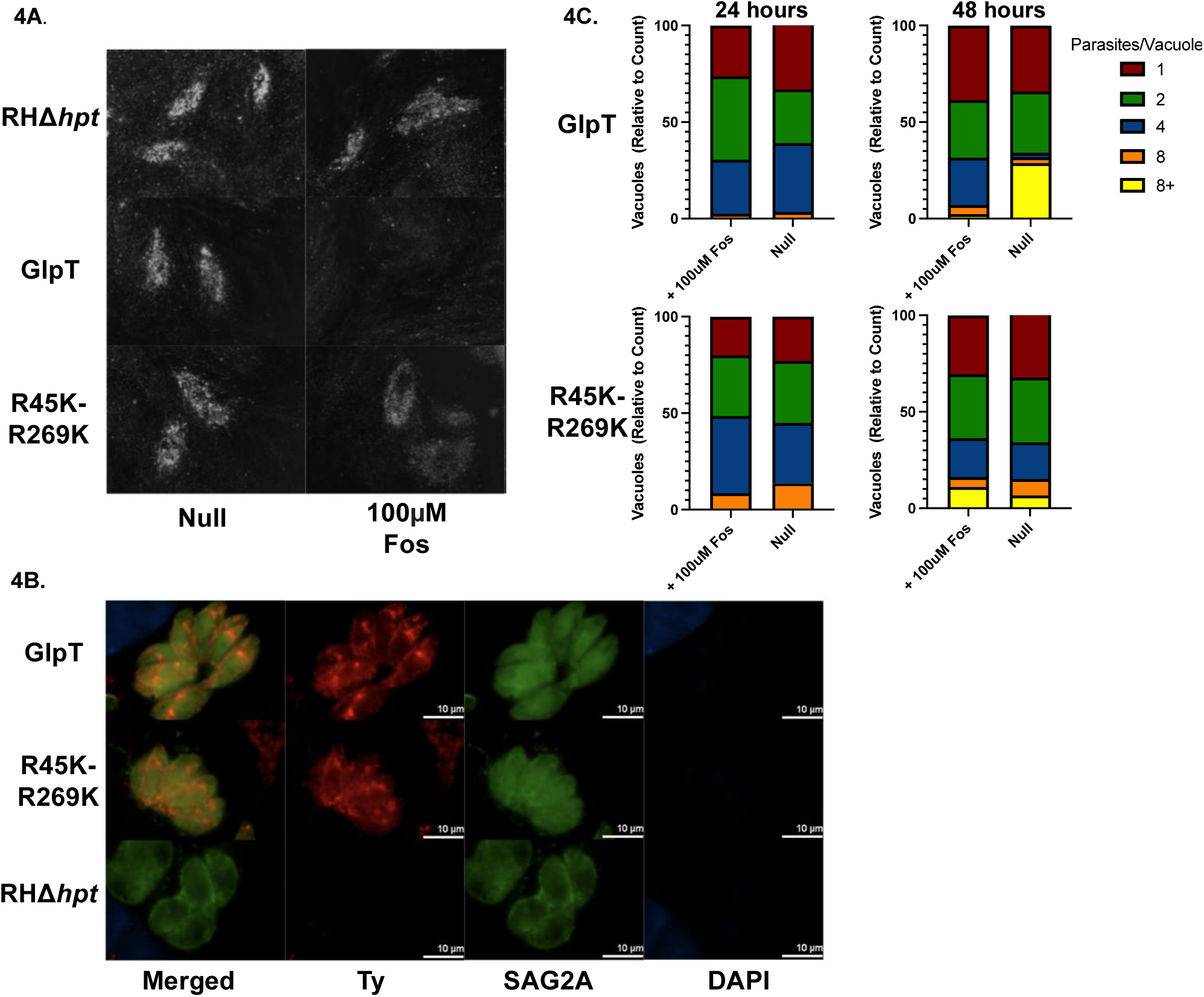
Fosmidomycin sensitivity in *Toxoplasma* tachyzoites requires expression of the GlpT transporter. (**A**) Plaque assays were performed using 24-well plates with HFFs pre-incubated for an hour with or without 100 µM fosmidomycin. Cultures were then infected with 300 tachyzoites of either the parental RHΔ*hpt* strain, the GlpT transgenic line (Ty-tagged GlpT, functional glycerol-3-phosphate transporter), or the point-mutated GlpT control (R45K-R269K). Plaques were inspected after 5 days. (**B**) Immunofluorescence assays confirmed expression and localization of GlpT in the GlpT and R45K-R269K strains. Staining was performed using mouse anti-Ty, rabbit anti-SAG2A (parasite surface marker), and DAPI. Images were acquired at 100x magnification. (**C**) Parasite replication was assessed by counting parasites per vacuole at 24 and 48 hours post-infection. HFF monolayers were pretreated with or without 100 µM fosmidomycin 1 hour prior to infection with GlpT or R45K-R269K parasites.

**Fig S5.**
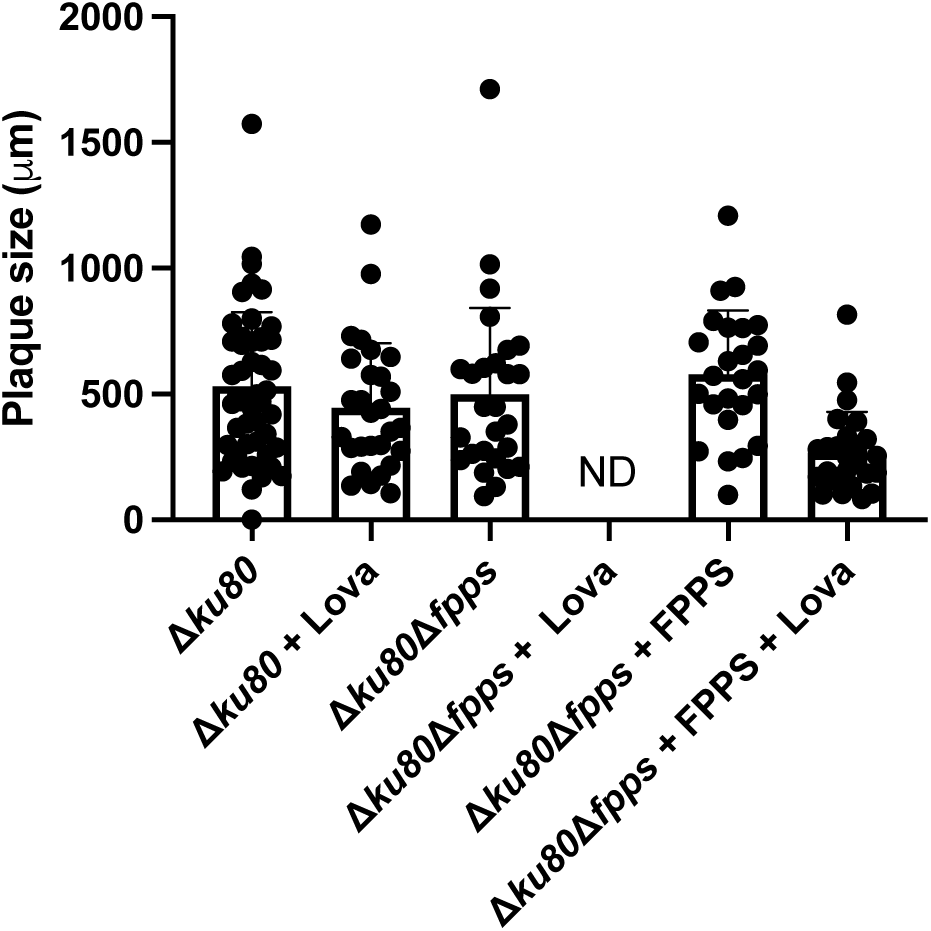
Lovastatin sensitivity confirms loss of FPPS activity in FPPS knockout parasites strains. Plaque assay was performed using 24-well plates with confluent HFF monolayers pretreated for 1 hour with or without 13 µM lovastatin. Cultures were then infected with 300 tachyzoites of RHΔ*ku80* (WT), RHΔ*ku80*Δfpps (FPPS knockout), or RHΔ*ku80*Δ*fpps* + FPPS (complemented). Cultures were then analyzed with confocal microscopy to measure plaque size. FPPS knockout parasites formed plaques in drug-free medium but failed to do so in lovastatin-treated wells. In contrast, WT and complemented parasites formed normal plaques under both conditions, confirming that sensitivity to lovastatin results from loss of endogenous FPPS. Plaques were not detected (ND) for the Δ*fpps* strain in the presence of lovastatin.

**Table S1.** Differential expression of host genes involved in the mevalonate pathway, phosphoantigen recognition, and lipid transport in PBMCs infected with *Toxoplasma gondii*. RNA-seq data from the GEO dataset GSE119835 were analyzed using GEO2R to compare gene expression between RH strain-infected and media-treated human PBMCs. In short, five PBMCs samples were split into three conditions: untreated, infected with RH88 (MOI 3), or infected with PRU (MOI 3). After 12 hours, RNA was extracted and sequenced (40). Differential expression was assessed by comparing RH-infected to untreated samples. Genes were selected based on known roles in the mevalonate/isoprenoid biosynthesis pathway, host phosphoantigen recognition, or lipid transport. Log2 fold change (log2FC) values represent expression differences (positive = upregulated; negative = downregulated). Adjusted p-values (Benjamini-Hochberg FDR) reflect the statistical significance of differentially expressed genes.

| Gene Symbol | Gene Name | log2FoldChange | Adjusted p-value | Pathway/Function |
| --- | --- | --- | --- | --- |
| SQLE | squalene epoxidase | 1.6 | 4.31E-05 | Cholesterol biosynthesis |
| HMGCS1 | 3-hydroxy-3-methylglutaryl-CoA synthase 1 | 1.3 | 1.7e-05 | Upstream mevalonate synthesis |
| MVK | mevalonate kinase | 1.2 | 2.0e-3 | Mevalonate phosphorylation |
| HMGCR | 3-hydroxy-3-methylglutaryl-CoA reductase | 1.1 | 3.4e-3 | Cholesterol biosynthesis |
| FDPS | farnesyl diphosphate synthase | 0.94 | 1.5e-04 | Isoprenoid synthesis |
| IDI1 | isopentenyl-diphosphate delta isomerase 1 | 1.0 | 1.6e-2 | IPP/DMAPP isomerization |
| MVD | mevalonate diphosphate decarboxylase | 0.95 | 1.7e-3 | Isoprenoid synthesis |
| PMVK | phosphomevalonate kinase | 0.15 | 6.9e-01 | Mevalonate phosphorylation |
| PPP2CA | protein phosphatase 2 catalytic subunit alpha | 0.48 | 1.9e-1 | PP2A catalytic subunit |
| ABCA1 | ATP binding cassette subfamily A member 1 | 0.075 | 9.2e-1 | Cholesterol efflux |
| IFNG | interferon gamma | 7.5 | 2.1e-26 | Cytokine |
| TNF | tumor necrosis factor | 1.7 | 5.1e-3 | Pro-inflammatory cytokine |
| IL12A | interleukin 12A | 0.92 | 7.0e-2 | Cytokine (IL-12 subunit) |
| IL1B | interleukin 1 beta | 3.8 | 2.1e-17 | Pro-inflammatory cytokine |
| BTN3A1 | butyrophilin subfamily 3 member A1 | -0.79 | 1.3e-1 | Phosphoantigen recognition (Vγ9Vδ2 T cell) |
| BTN2A1 | butyrophilin subfamily 2 member A1 | -0.42 | 3.0e-01 | BTN2 family |
| IL18 | interleukin 18 | -0.62 | 4.8e-01 | Inflammasome-related cytokine |
| GGPS1 | geranylgeranyl diphosphate synthase 1 | -0.55 | 1.5E-02 | Isoprenoid synthesis |

**Table S2.**
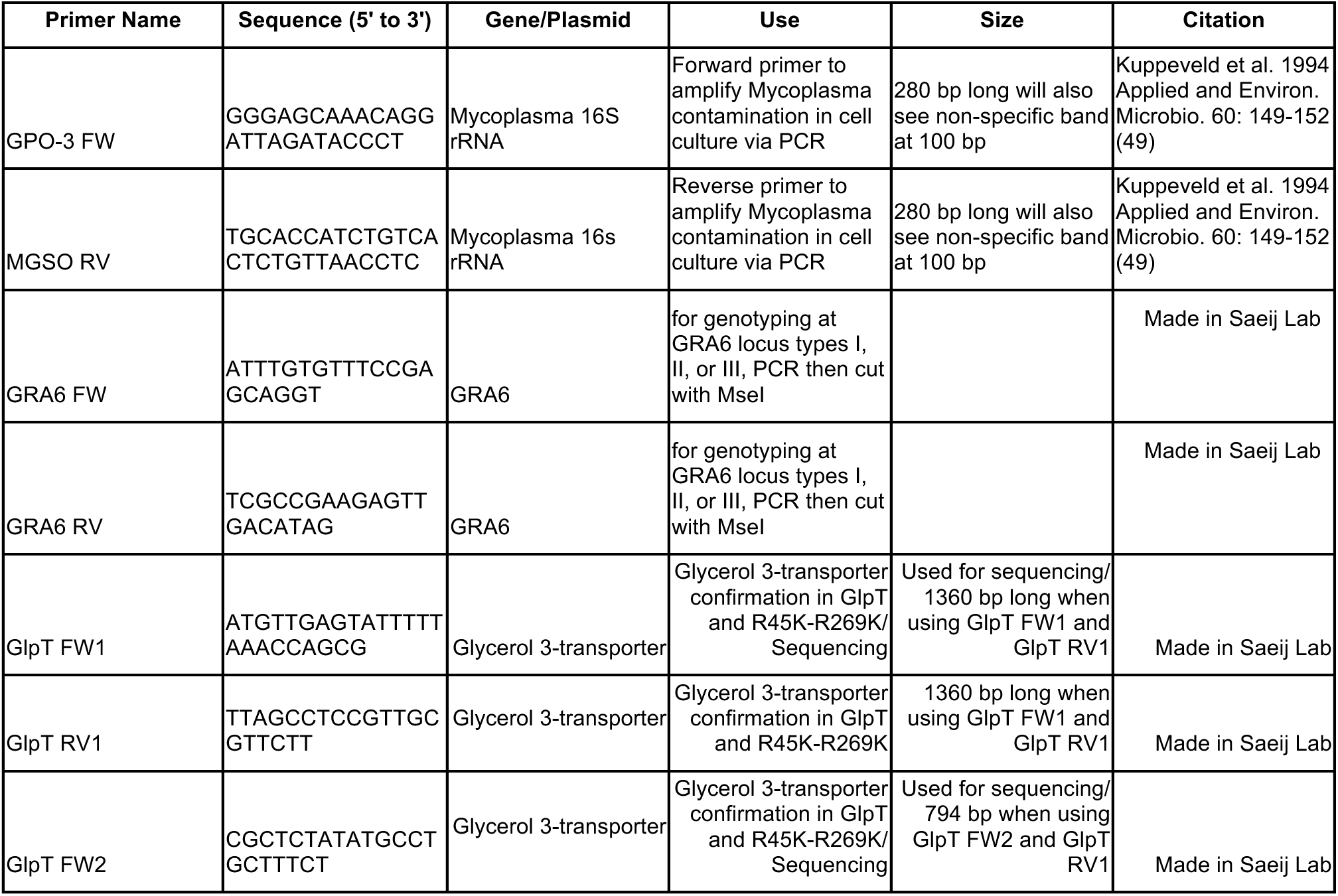
Primers used to test for mycoplasma, glycerol 3-transporter, and strain type.

